# A novel protocol for magnetoencephalographic mapping of speech motor control regions during speech production

**DOI:** 10.1101/2023.09.30.560288

**Authors:** Ioanna Anastasopoulou, Douglas O. Cheyne, Pascal van Lieshout, Kirrie Ballard, Peter Wilson, Blake W Johnson

## Abstract

Neuroimaging protocols for mapping of expressive speech centres employ several standard speech tasks including object naming, rhyming, and covert word production (Agarwal et al., 2019). These tasks reliably elicit activation of distributed speech centres in prefrontal, precentral and cingulate motor cortices and are widely used for presurgical mapping and in research studies of language production. In the present study we used an alternative speech protocol employing reiterated productions of simple disyllabic nonwords (VCV; Anastasopoulou et al., 2022; van Lieshout et al., 2007). Here we show that this task elicits highly focal and highly lateralised activations of speech motor control areas centred on the precentral gyrus and adjacent portions of the middle frontal gyrus. 10 healthy adults, 19 typically developing children and 7 children with CAS participated in the study. MEG scans were carried out with a whole-head MEG system consisting of 160 first-order axial gradiometers with a 50 mm baseline (Model PQ1160R-N2, KIT, Kanazawa, Japan). MEG data were acquired with analog filter settings of 0.03 Hz high-pass, 1,000 Hz low-pass, 4,000 Hz sampling rate. Measurements were carried out with participants in supine position in a magnetically shielded room (Fujihara Co. Ltd., Tokyo, Japan). Time-aligned speech acoustics were recorded in an auxiliary channel of the MEG setup at the same sample rate as the MEG recordings. Brain activity was recorded while participants produced reiterated utterances of /ipa/ and /api/, at normal and speeded rates in addition to a button press task (right index finger) to elicit activity in the hand region of sensorimotor cortex (e.g. Johnson et al., 2020). MEG data were co-registered with individual structural MRI scans obtained in a separate scanning session. Source reconstruction was performed with synthetic aperture magnetometry (SAM) beamformer implemented in Matlab (Jobst et al., 2018) and group statistics performed with permutation tests (p < 0.05). Button press map shows clusters encompassing dorsal precentral and postcentral gyri (Brodmann areas 4 and 6), corresponding to hand sensorimotor cortices. Speech map shows clusters encompassing precentral gyrus immediately ventral to hand motor cortex (BA6), and an immediately adjacent portion of the posterior middle frontal gyrus. Both button press and speech result in a robust desynchronisation restricted within a frequency band of about 13-30 Hz (beta band). Our results show that the reiterated speech task results in robust beta-band desynchronisation in a highly focal region of the precentral gyrus, located immediately ventral to the hand motor region of the precentral gyrus. In adults, speech motor -related brain activity was predominantly observed in the left hemisphere. Typically developing children, on the other hand, exhibited bilateral activation and in the case of individuals with CAS exhibited only right-hemisphere activation. Taken together the present findings provide a non-invasive and highly selective window on a crucial node of the expressive speech network that has previously been accessed only with invasive electrophysiological means and lesion studies.

## Introduction

Since the advent of modern functional neuroimaging techniques, an important clinical and scientific aim has been to develop and refine methodologies for imaging brain function during overt speaking. From a basic science perspective these methods are integral to achieving an understanding of the brain regions and networks that interact with speech comprehension systems, generate speech plans, and ultimately send movement commands to some 100 muscles associated with the articulators of the vocal tract. In a biomedical context expressive speech mapping methods are an important component of neurosurgical planning where any potential for damage to brain speech centres would inevitably have devastating and irreversible consequences for the postoperative communicative and cognitive capabilities of the patient.

Currently, two main neuroimaging techniques are used for non-invasive mapping of expressive speech function. Functional magnetic resonance imaging (fMRI) relies on now fairly ubiquitous MRI scanning technology and is dominant to a considerable degree in both clinical and experimental contexts. Here our focus is on magnetoencephalography (MEG), a more specialised technology that relies on less widely available scanning equipment. Its use for expressive language mapping is accordingly restricted to the limited number of clinical sites and laboratories with access to an MEG scanner: there are currently about 20 of these sites in the United States, and several hundred worldwide.

Progress in expressive language and other types of brain mapping will likely continue to require both scanning techniques as they have well-known, distinctive, and in many ways complementary advantages and disadvantages. Notably, fMRI relies on an indirect measure of neuronal activity (the hemodynamic response to neuronal metabolism) which is relatively spatially precise but temporally sluggish. Conversely, MEG has direct access to the magnetic fields generated by neuronal activities. Neuromagnetic fields accordingly provide high temporal resolution for brain activity, an important consideration in the context of a rapid and dynamic behaviour like speech; however the spatial specification (localisation) of electromagnetic fields is a physically complex problem, and in practice spatial resolution can vary from completely ambiguous to sub-centimetre precision, depending largely on how extensive and complex are the configurations of active neural generators in a given behaviour or task. Speech is a complex behaviour that draws on multiple brain centres and systems, and hence in clinical contexts MEG expressive speech mapping precision is often limited to an assessment of hemispheric lateralisation, rather than for a more detailed specification of any brain region(s) within hemispheres.

A fairly extensive array of well-defined speaking tasks/protocols are now available for MEG mapping of expressive brain function, the selection of which depends on clinical and laboratory experience and preference as well as the specific clinical or experimental aims of a given neuroimaging session (see Agarwal, 2019 for a review of comparable speech protocols used for fMRI expressive language mapping). These tasks include, in rough descending order of the overall linguistic/cognitive task demands: sentence reading; semantic word judgements (e.g. abstract/concrete); word recognition; picture naming; verb generation; word reading (single words, phrases, or lists); nonword repetition; and non-linguistic oromotor gestures. See Appendix 1 for a list of MEG studies (organised by speech task) of expressive language function published since 2015; see also Munding et al. (2015) for a review and list of MEG expressive language studies published prior to 2015). There is considerable overlap in the brain regions activated by different speech tasks (Agarwal et al., 2019), but in general it can be stated that tasks that require access to high level memorial and cognitive/linguistic operations tend to activate distributed areas of prefrontal, temporal and parietal cortex, including Broca’s area in the left hemisphere (Bowyer et al., 2005; Doesburg et al., 2016; Kadis et al., 2011; Youssofzadeh & Babajani-Feremi, 2019; Correia et al., 2020). At the other end of the spectrum of task complexity, non-word/pseudoword and oromotor tasks are intended to limit the requirements for semantic, syntactic and attentional processing and elicit neural activity that is more restricted to brain regions associated with phonological, phonetic and sensorimotor processes (Frankford et al., 2021).

The present MEG study was designed to assess speech motor cortex activations elicited by a *reiterated non-word speech production task* that is commonly used to investigate speech motor control by behavioural means but that has rarely been used in neuroimaging studies of expressive language. To our knowledge, only one early fMRI study has explicitly investigated expressive speech mapping using reiterated non-lexical speech. Riecker et al. (2000) measured brain activity with fMRI from healthy, right-handed native German speakers. Participants were required to produce monosyllables (“ta” and “stra”), a non-lexical syllable sequence (“pataka”), a lexical item (“Tagebau”) and horizontal tongue movements. Participants were instructed to produce the test items in a monotonous manner at a self-paced comfortable speaking rate during measurement periods extending to 1 min, and to refrain from covert verbalisation during the inter-trial rest periods. These authors reported that all speech-rest contrasts resulted in significant activations restricted to the ventral portion of the peri-Rolandic sensorimotor cortices. Bilateral activations were found for tongue movements, “ta”, “stra” and “Tagebau”, while “pataka” showed only left hemispheric activation. The authors concluded that that the highly focal and restricted neural activations associated with reiterated speech may reflect the “chunking” or organisation of coarticulated syllable strings into a single output unit, which places fewer demands on neural resources relative to production of smaller or single (“individualised”) units of speech or action.

As noted above, MEG source reconstruction is more amenable to restricted and focal than to widespread and distributed source configurations; it follows that a speech task that favours the former configurations will provide a more precise and informative mapping of expressive language function than existing protocols that remain largely limited to a specification of hemispheric laterality. In the following, we describe our MEG results for expressive language mapping using reiterated nonword tasks in healthy adults, and contrast these with mapping results from typically developing children. We also describe results from a group of children with Childhood Apraxia of Speech (CAS), a developmental motor speech disorder that is believed to result from a central deficit in the ability to program the syllable sequences and transitions that are required for successful performance of reiterated speech tasks.

## Methods

### Participants

Three groups were recruited for this study. Eleven healthy adults (4 females, mean age = 35.5 years; Standard Deviation [SD] = 15.0, range 19.8 – 64.6), 19 typically developing (TD) children (8 females, mean age =11.0 years; SD = 2.5, range 7.5 – 16.7) and seven children with Childhood Apraxia of Speech (CAS) (7 males, mean age = 8.9 years, SD = 2.2, range = 6.8 - 12.8) (for detailed demographics see Appendix 3). The typically developing children were distributed into two distinct age groups to investigate potential developmental disparities in language localization. The younger group (n = 9) encompassed children whose ages ranged from 7.45 to 9.55 years and the older group (n = 10) included participants aged from 11.06 to 16.73 years. The utilization of these age-defined subgroups facilitated the examination of nuanced differences in language localization capabilities across varying stages of development. All procedures were approved by the Macquarie University Human Subjects Research Ethics Committee.

### Speech and motor assessments

All children passed a pure-tone hearing screening for the frequencies of 1000, 2000, and 4000 Hz at 20 dB and 500 Hz at 25 dB. All child participants completed a battery of speech, expressive and receptive language, and motor assessments: The Sounds-in-Words subtest of the Goldman-Fristoe Test of Articulation–Third Edition (GFTA-3; Goldman & Fristoe, 2015); the Receptive and Expressive Language components of the Clinical Evaluation of Language Fundamentals Fifth Edition (CELF-5; Wiig, Secord, & Semel, 2013); nonverbal components of the Reynolds Intellectual Assessment Scales (Reynolds & Kamphaus, 2003); Verbal Motor Production Assessment for Children (VMPAC-R, Hayden, & Namasivayam, 2021); the Single-Word Test of Polysyllables (Gozzard, Baker, & McCabe, 2004, 2008). This test consists of a 50-item picture-naming task designed to assess articulation, sound, and syllable sequencing, as well as lexical stress accuracy (i.e., prosody) (Murray et al., 2015), and the Movement Assessment Battery for Children Second Edition (ABC-2, Henderson et al., 2007). Further, caregivers completed two questionnaires: the Developmental Coordination Disorder Questionnaire (DCDQ, Wilson et al.,2009) and the Handedness Questionnaire (HQ, Oldfield, 1971, Veale, 2014).

### CAS diagnosis and group assignment

Initially, ten children with suspected CAS were enrolled in this study. Two certified speech pathologists, authors I.A. and K.B, independently reviewed assessment videos of all ten children with CAS. They assessed each child based on their perceptual evaluation of their speech samples, using the following procedure. In order to receive a CAS diagnosis in our study, participants needed to exhibit (a) the three features established by consensus in the ASHA Technical Report (2007b) and (b) a minimum of four out of the 10 features outlined in Strand’s 10-point checklist (Shriberg, Potter, et al., 2009, Murray et al., 2015) across at least three assessment tasks, as detailed in Table 1.

**Table 1.** Assessment tasks and diagnostic features used for assigning an expert diagnosis of Childhood Apraxia of Speech and Developmental Coordination Disorder.

| CAS |  |  |
| --- | --- | --- |
| ASHA consensus-based feature list | Strand's 10-point checklist | Assessment tasks taken into account for the diagnosis |
| Inconsistent errors on consonants and vowels in repeated productions of syllables or words | Difficulty achieving initial articulatory configurations and transitions into vowels (within-speech groping, false starts, restarts, and hesitations) | Single-Word Test of Polysyllables (Gozzard, Baker, & McCabe, 2004, 2008) |
| Lengthened and disrupted coarticulatory transitions between sounds and syllables | Syllable segregation | Diadochokinetic task /pataka/ from the VMPAC assessment (Hayden, & Namasivayam, 2021) |
| Inappropriate prosody, especially in the realization of lexical or phrasal | Lexical stress errors or equal stress | GFTA Sentences (GFTA-3; Goldman & Fristoe, 2015) |
|  | Vowel or consonant distortions including distorted substitutions | CELF Sentences (CELF-5; Wiig, Secord, & Semel, 2013) |
|  | Groping (nonspeech, oral groping) |  |
|  | Intrusive schwa |  |
|  | Voicing errors |  |
|  | Slow rate |  |
|  | Increased difficulty with longer or more phonetically complex words |  |

Table 2 displays the assessment results for the presence or absence of Childhood Apraxia of Speech (CAS) features in each of the 10 participants. Out of these 10 individuals, 8 participants (80%) were diagnosed with CAS, as per the criteria outlined by ASHA (2007b) and Shriberg, Potter, et al. (2009). Of the original ten children in the cohort, two did not display any features associated with apraxia of speech. Additionally, one child in this cohort was left-handed and therefore did not participate in the study. The remaining seven children all took part in the study, and the interrater reliability was determined to be 87.5%.

**Table 2.**
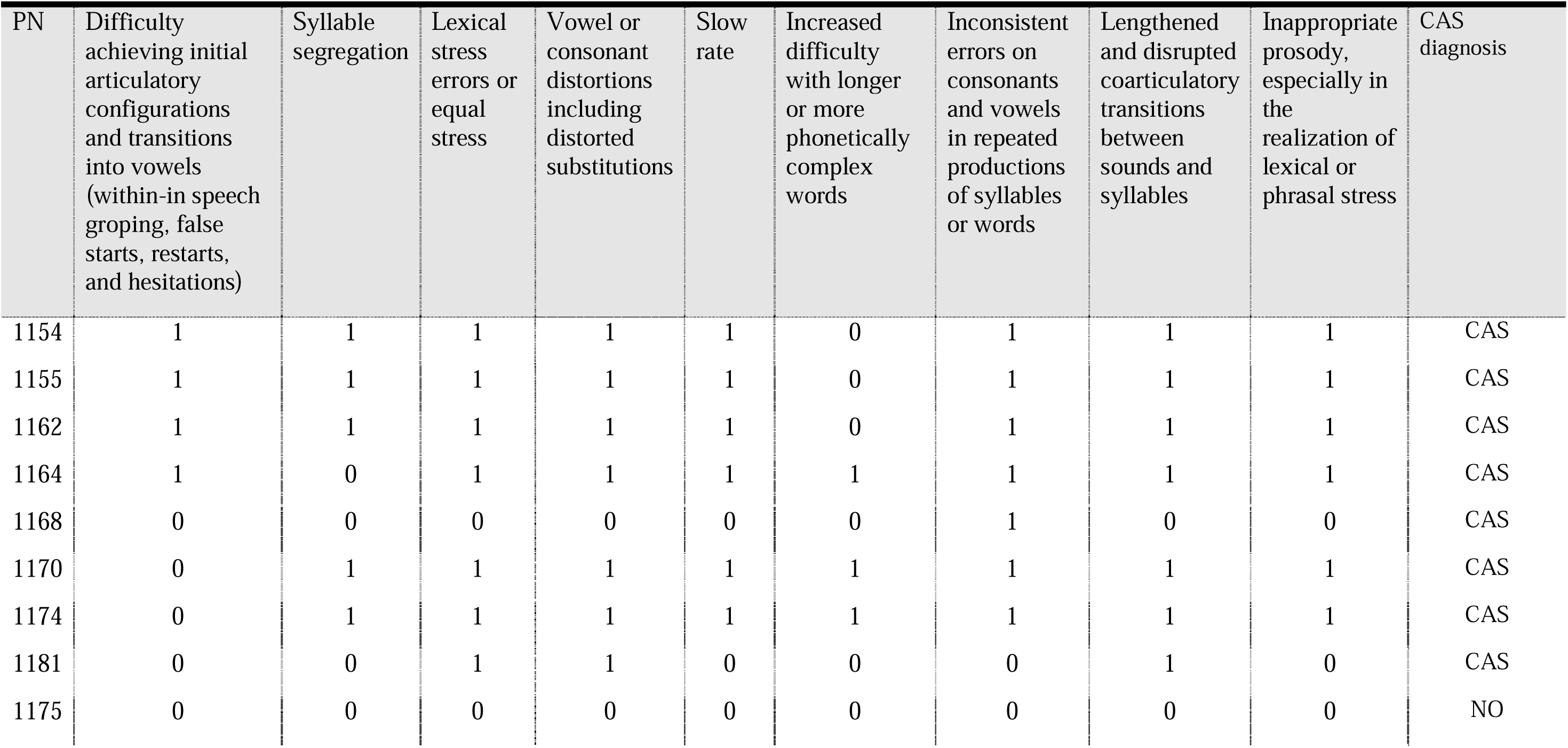
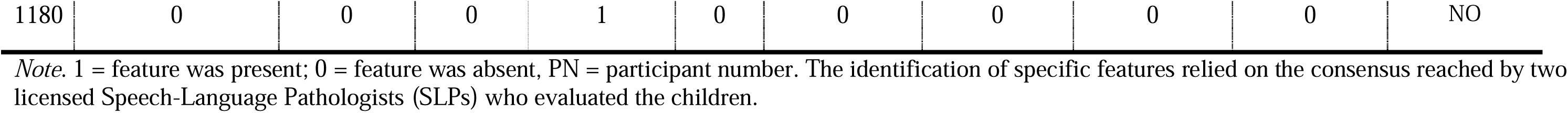
Presence of childhood apraxia of speech (CAS) features for individual participants.

| PN | Difficulty achieving initial articulatory configurations and transitions into vowels (within-in speech groping, false starts, restarts, and hesitations) | Syllable segregation | Lexical stress errors or equal stress | Vowel or consonant distortions including distorted substitutions | Slow rate | Increased difficulty with longer or more phonetically complex words | Inconsistent errors on consonants and vowels in repeated productions of syllables or words | Lengthened and disrupted coarticulatory transitions between sounds and syllables | Inappropriate prosody, especially in the realization of lexical or phrasal stress | CAS diagnosis |
| --- | --- | --- | --- | --- | --- | --- | --- | --- | --- | --- |
| 1154 | 1 | 1 | 1 | 1 | 1 | 0 | 1 | 1 | 1 | CAS |
| 1155 | 1 | 1 | 1 | 1 | 1 | 0 | 1 | 1 | 1 | CAS |
| 1162 | 1 | 1 | 1 | 1 | 1 | 0 | 1 | 1 | 1 | CAS |
| 1164 | 1 | 0 | 1 | 1 | 1 | 1 | 1 | 1 | 1 | CAS |
| 1168 | 0 | 0 | 0 | 0 | 0 | 0 | 1 | 0 | 0 | CAS |
| 1170 | 0 | 1 | 1 | 1 | 1 | 1 | 1 | 1 | 1 | CAS |
| 1174 | 0 | 1 | 1 | 1 | 1 | 1 | 1 | 1 | 1 | CAS |
| 1181 | 0 | 0 | 1 | 1 | 0 | 0 | 0 | 1 | 0 | CAS |
| 1175 | 0 | 0 | 0 | 0 | 0 | 0 | 0 | 0 | 0 | NO |
| Expressive speech mapping with MEG |  |  |  |  |  |  |  |  |  |  |
| 1180 | 0 | 0 | 0 | 1 | 0 | 0 | 0 | 0 | 0 | NO |

*Note.* 1 = feature was present; 0 = feature was absent, PN = participant number. The identification of specific features relied on the consensus reached by two licensed Speech-Language Pathologists (SLPs) who evaluated the children.

Descriptive statistics comparing three groups, namely a group of 19 children with typical development referred to as the TD group, a group of 7 children diagnosed with Childhood Apraxia of Speech (CAS) referred to as the CAS group, and 7 age - matched controls selected from the TD cohort, are presented in Table 3. Notably, no statistically significant differences were observed between the CAS group and the age-matched control group in terms of age, nonverbal IQ, as well as expressive and receptive language abilities as measured by the CELF-5 assessment (Wiig, Secord, & Semel, 2013).

**Table 3.**
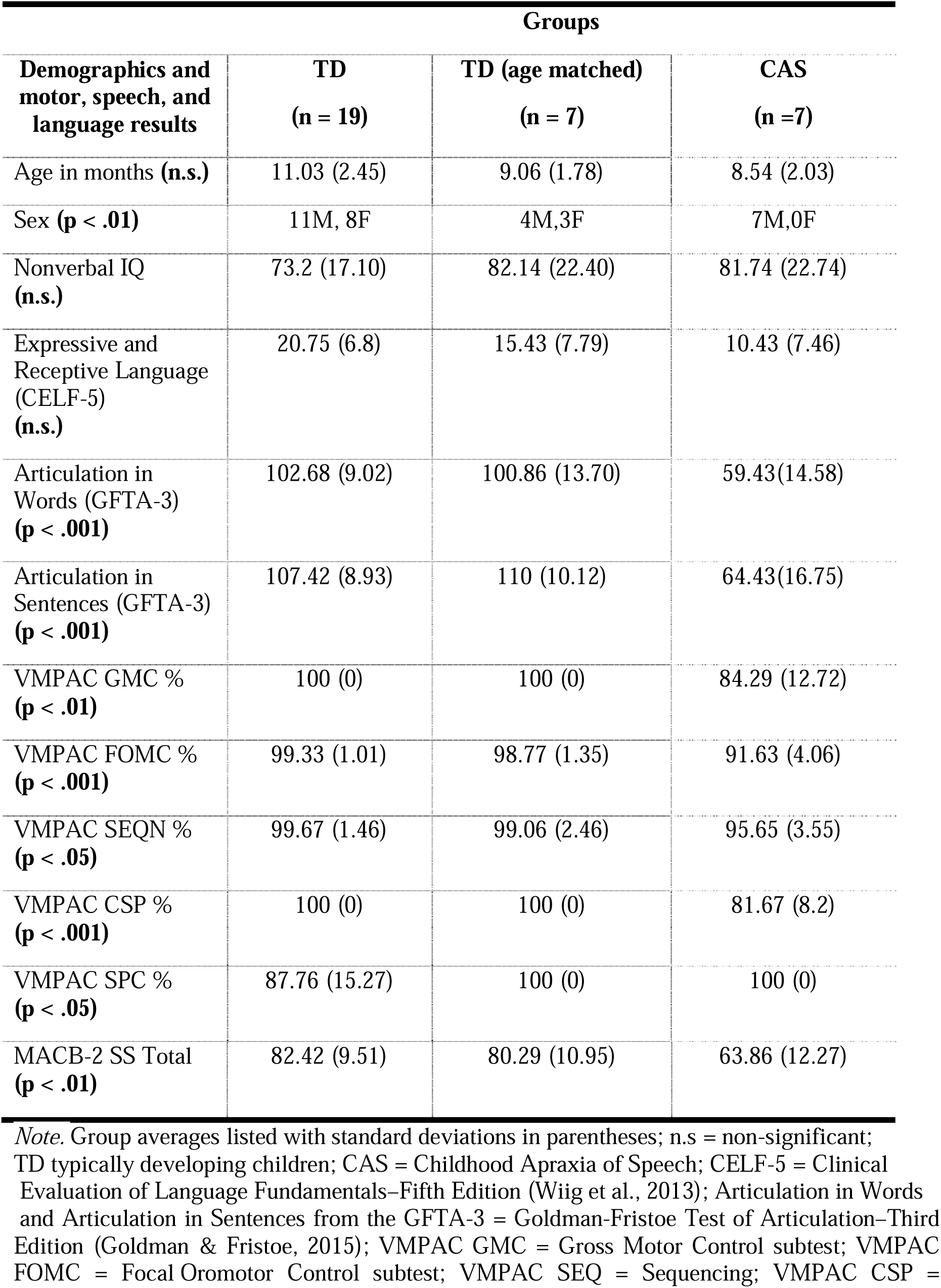
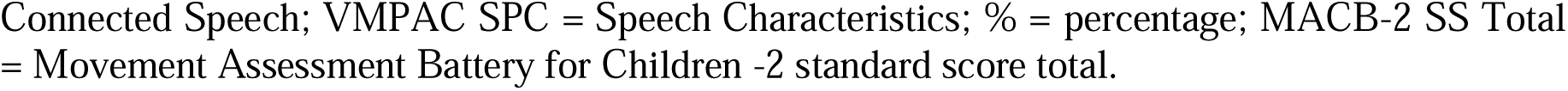
Demographics and motor, speech, and language results by group.

| **Demographics and motor, speech, and language results** | **Groups** |  |  |
| --- | --- | --- | --- |
|  | **TD**<br>**(n = 19)** | **TD (age matched)**<br>**(n = 7)** | **CAS**<br>**(n = 7)** |
| Age in months (**n.s.**) | 11.03 (2.45) | 9.06 (1.78) | 8.54 (2.03) |
| Sex (**p < .01**) | 11M, 8F | 4M,3F | 7M,0F |
| Nonverbal IQ (**n.s.**) | 73.2 (17.10) | 82.14 (22.40) | 81.74 (22.74) |
| Expressive and Receptive Language (CELF-5) (**n.s.**) | 20.75 (6.8) | 15.43 (7.79) | 10.43 (7.46) |
| Articulation in Words (GFTA-3) (**p < .001**) | 102.68 (9.02) | 100.86 (13.70) | 59.43(14.58) |
| Articulation in Sentences (GFTA-3) (**p < .001**) | 107.42 (8.93) | 110 (10.12) | 64.43(16.75) |
| VMPAC GMC % (**p < .01**) | 100 (0) | 100 (0) | 84.29 (12.72) |
| VMPAC FOMC % (**p < .001**) | 99.33 (1.01) | 98.77 (1.35) | 91.63 (4.06) |
| VMPAC SEQN % (**p < .05**) | 99.67 (1.46) | 99.06 (2.46) | 95.65 (3.55) |
| VMPAC CSP % (**p < .001**) | 100 (0) | 100 (0) | 81.67 (8.2) |
| VMPAC SPC % (**p < .05**) | 87.76 (15.27) | 100 (0) | 100 (0) |
| MACB-2 SS Total (**p < .01**) | 82.42 (9.51) | 80.29 (10.95) | 63.86 (12.27) |

*Note.* Group averages listed with standard deviations in parentheses; n.s = non-significant; TD typically developing children; CAS = Childhood Apraxia of Speech; CELF-5 = Clinical Evaluation of Language Fundamentals–Fifth Edition (Wiig et al., 2013); Articulation in Words and Articulation in Sentences from the GFTA-3 = Goldman-Fristoe Test of Articulation–Third Edition (Goldman & Fristoe, 2015); VMPAC GMC = Gross Motor Control subtest; VMPAC FOMC = Focal Oromotor Control subtest; VMPAC SEQ = Sequencing; VMPAC CSP =
Connected Speech; VMPAC SPC = Speech Characteristics; % = percentage; MACB-2 SS Total = Movement Assessment Battery for Children -2 standard score total.

As expected, statistically significant differences were obtained for articulation skills assessment, encompassing both word and sentence contexts as evaluated by the Goldman-Fristoe Test of Articulation-3 (GFTA-3; Goldman & Fristoe, 2015) across the two groups. In addition, all constituent components of the VMPAC (Hayden & Namasivayam, 2021) yielded statistically significant findings.

Furthermore, outcomes derived from the Movement Assessment Battery for Children Second Edition (MABC-2, Henderson et al., 2007) demonstrated statistically significant distinctions between the cohorts, suggestive of the presence of features associated with Developmental Coordination Disorder (DCD) in children with CAS.

### MEG acquisition

Neuromagnetic brain activity was recorded with a KIT-Macquarie MEG160 (Model PQ1160R-N2, KIT, Kanazawa, Japan) whole-head MEG system consisting of 160 first-order axial gradiometers with a 50-mm baseline (Kado et al., 1999; Uehara et al., 2003). MEG data were acquired with analogue filter settings as 0.3 Hz high-pass, 200 Hz low-pass, 1000 Hz sampling rate and 16-bit quantization precision. Measurements were carried out with participants in supine position in a magnetically shielded room (Fujihara Co. Ltd., Tokyo, Japan). Five head position indicator coils (HPI) were attached in the head in an elastic cap, and their positions were measured at the beginning and at the end of the experiment, with a maximum displacement criterion of < 5 mm in any direction. The coils’ positions with respect to the three anatomical landmarks (nasion, right and left preauricular landmarks) were measured using a handheld digitiser (Polhemus FastTrack; Colchester, VT).

Participant’s head shapes and fiducial positions were digitized using the Polhemus FastTrack system (Polhemus FastTrack; Colchester, VT), enabling later co-registration with anatomical MRIs (Gross et al., 2013; Mersov et al., 2016). To quantify head movement, marker coil positions affixed to an elastic cap were measured twice; at the beginning and the end of the session; with a maximum displacement criterion of less than 5 mm in any direction.

T1-weighted anatomical magnetic resonance images (MRIs) were acquired for all adult participants in a separate scanning session using a 3T Siemens Magnetom Verio scanner with a 12-channel head coil. Those anatomical images were obtained using 3D GR\IR scanning sequence with the following acquisition parameters: repetition time, 2000 ms; echo time, 3.94 ms; flip angle, 9 degrees; slice thickness, 0.93 mm; field of view, 240 mm; image dimensions, 512 × 512 × 208. For child participants we used a “surrogate” MRI approach which warps a template brain to each subject’s digitized head shape using the iterative closest point algorithm implemented in SPM8 (Litvak et al., 2011) and the template scalp surface extracted with the FSL toolbox (Jenkinson et al., 2012; see Cheyne et al., 2014; Johnson et al., 2022). Time-aligned audio speech recordings were recorded in an auxiliary channel of the MEG setup with the same sample rate (1000 Hz) as the MEG recordings.

### Speech movement tracking and high-fidelity acoustic recordings

All participants were fitted with MEG-compatible speech movement tracking coils place on the upper and lower lips, tongue body, and jaw (Alves et al., 2016; Anastasopoulou et al., 2022). Further, high fidelity speech recordings were simultaneously recorded with an optical microphone (Optoacoustics, Or-Yehuda, Israel) fixed on the MEG dewar at a distance of 20 cm away from the mouth of the speaker; and digitised using a Creative sound blaster X-Fi Titanium HD sound card with 48 kHz sample rate and 24-bit quantization precision.

The analyses of the present paper utilise only speech onset/offset events identified from the acoustic recordings of the MEG auxiliary channel. Analyses of the speech tracking and high-fidelity acoustic data are presented in Chapter 5 (and Anastasopoulou et al., 2023 preprint) and are not further discussed here.

### Experimental protocol

Four non-word productions were used as experimental stimuli, consisting of two disyllabic sequences with a V1CV2 structure, /ipa/ and /api/, each produced at normal and faster rates. These stimuli have been used in previous studies to investigate speech motor control strategies in both normal and disordered populations investigating speech motor control strategies in normal and in disordered populations (van Lieshout et al. 1997; van Lieshout et al. 2002; van Lieshout et al. 2007; van Lieshout, 2017). Non-word stimuli were chosen to avoid familiarity issues and are commonly used in speech motor control research to investigate normal and pathological function (Case & Grigos, 2020). The general experimental protocol for speech and button press conditions is diagrammed in Figure 1. Each participant repeated all tasks in ten trials, with each trial lasting approximately 12 seconds. They were instructed to take a deep breath and, for the normal rate production, to utter the non-words in a comfortable, conversational rate. For the faster rate, they were instructed to produce the stimuli at a faster rate while maintaining accuracy (van Lieshout et al., 2002). A short break was provided after each trial. Participants were asked to minimize head movement as much as possible and avoid blinking their eyes during speech production (Mersov et al., 2016).

**Figure 1.**
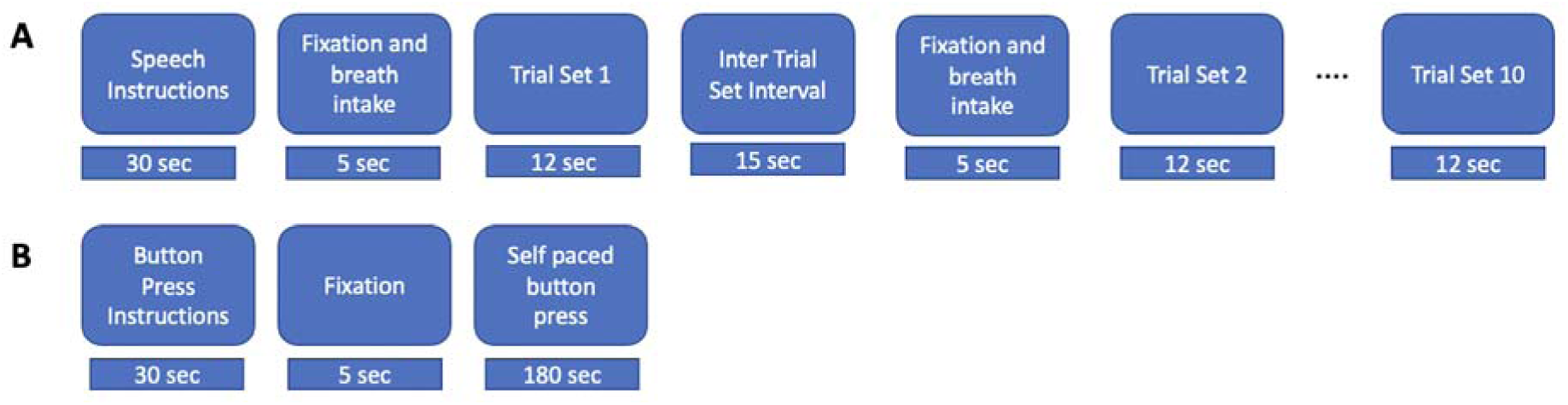
Experimental procedures. *Note.* **A.** Speech task. Instructions were displayed for 30s, followed by a 5s fixation cross ‘+’ and breath intake in preparation for the speech production trial set. During a trial set participants produced the indicated nonword in a reiterated fashion for 12s. 10 consecutive trial sets were performed for each nonword stimulus. **B.** Button press task. Instructions were displayed for about 30s followed by a fixation cross, during which participants performed self-paced button pressed with the index finger of their dominant (right) hand at a rate of about 1 per 2 seconds for a total of about 90 trials.

## Analysis

### Button press condition

MEG data were segmented into 1.5 s epochs comprising -0.5 sec to +1.0 sec with respect to button press onset, and about 90 trials per averaged epoch. Epoched data sets were digitally filtered from 0.3-100 Hz and a 50 Hz notch filter. SAM pseudo-t beamformer was computed using a frequency range of 15-25 Hz (beta band), a sliding active window of 0.6 to 0.8 sec and a baseline window of 0.0 to 0.2 sec (first two seconds of inter trial set rest period) over 10 steps with a step size of 0.01 sec and pseudo-z beamformer normalisation (3 ft/sqrt (Hz) RMS noise). Active and baseline windows were chosen to encompass the known times of maximal event-related synchronisation (ERS) and event-related desynchronisation (ERD) respectively of MEG motor rhythms in a button press task (see for e.g., Cheyne et al., 2014; and Johnson et al., 2022). Statistical analysis of group beamformer images was performed with cluster-based permutation testing (2048 permutations, omnibus correction for multiple comparisons). Voxel locations at the centre of significant clusters were used to generate group mean “virtual sensor” time frequency plots with a time range of -0.5 to +1.0 sec from the button press onset and a frequency range of 1-100 Hz.

### Speech conditions

Speech trial set onsets were identified and marked from the speech channel of the MEG recordings. MEG data were segmented with an epoch of -10 sec to + 5 sec from the onset of each trial set, selected to encompass the final 5 sec of the previous trial set, the 5 sec inter-trial set rest period, and the first five sec of the current trial set (speech – rest – speech). To maximize the signal-to-noise ratio of the averaged data, all four speech conditions were averaged, for a total of 40 trial sets/averaged epoch (4 speech conditions * 10 speaking/resting trial sets). All epoched data were digitally filtered with a bandpass of 0.3-100 Hz and a 50 Hz notch filter.

Source reconstruction was performed using the scalar synthetic aperture magnetometry (SAM) beamformer implemented in the BrainWave MATLAB toolbox (Jobst et al., 2018). SAM pseudo-t beamformer was computed using a frequency range of 15-25 Hz, a sliding active window of 0 to 1.0 sec (first second of current speech trial set) and a baseline window of -5 to -3 sec (first two seconds of inter trial set rest period) over 10 steps with a step size of 0.2 sec and pseudo-z beamformer normalisation (3 ft/sqrt (Hz) RMS noise). Statistical analysis of group beamformer images was performed with cluster-based permutation testing (alpha = 0.05, 512-1024 permutations, omnibus correction for multiple comparisons). Voxel locations at the centre of significant clusters were used to generate group mean “virtual sensor” time frequency plots with a time range of -10 to +5 sec from the trial set onset and a frequency range of 1-100 Hz.

## Results

### Button press data

Group statistical analyses of SAM-beamformer images for the button-press condition are summarised in Table 4, and the corresponding group mean maps are shown in Figure 3. As expected, all groups show a single significant cluster located near the hand region of the left precentral gyrus, contralateral to the right-handed button press. As is typically observed for button press tasks, all groups also showed mirrored right hemisphere activations in homologous regions of the right hemisphere motor cortex (not shown), although these clusters were smaller in magnitude and did not reach statistical significance for any of the groups. Relative to adults, the cluster centre *Z* coordinate locations (superior-inferior direction) for the TD groups were 23 mm inferior, while the CAS group was 12 mm inferior to the adults. This discrepancy is largely due to the fact that the precise voxel maximum for the fairly extensive clusters of the adult data was located superior to the anatomic location of the hand knob. For the TD groups, the group mean maps of Figure 3 suggests that activations are somewhat deeper and located in the sulci anterior and posterior to the precentral gyrus, relative to the maps for adults and CAS children, which show prominent activations on the crests of the pre- and post-central gyri: however the present analyses do not permit inferences concerning whether these are true group differences (Sassenhagen and Drashkow, 2019).

**Table 4.** Centre coordinates of significant clusters for button press condition from cluster-based permutation analysis of SAM-beamformer source maps.

| Tal Coordinates |  |  |  |  |  |
| --- | --- | --- | --- | --- | --- |
| Group | X | Y | Z | Mag (nAm) | Brain Region |
| Adults | -34 | -13 | 66 | 6.0 | L Precentral Gyrus BA4 |
| TD 11-16 yo | -30 | -17 | 43 | 7.0 | L Precentral Gyrus BA4 |
| TD 7-10 yo | -34 | -16 | 43 | 6.6 | L Precentral Gyrus BA4 |
| CAS | -34 | -21 | 54 | 6.5 | L Precentral Gyrus BA4 |

*Note.* Cluster alpha = 0.05, 512-1024 permutations, omnibus correction for multiple comparisons. Locations are in Talairach coordinate system. L = left hemisphere. Group mean maps are shown in Figure 3.

Group mean virtual sensor time-frequency plots were generated from the button press coordinates listed in Table 4 and are plotted in the left panel of Figure 4. The following time-frequency features can be observed in the plots for all four groups: (1) A short discrete burst of gamma-band (circa 60-80+ Hz) synchronisation at or shortly before the button press; (2) Beta-band (circa 13-30 Hz) desynchronisation beginning about 300 ms before the button press and persisting for about 300-400 ms after the button press; (4) Beta-band synchronisation (beta “rebound”) beginning about 500-600 ms after the button press and persisting until the end of the analysis epoch; theta-band (circa 3-7 Hz) synchronisation beginning about 200-300 ms prior to the button press and persisting until 500-600 ms post-button press. All of these temporal-spectral features are known characteristics of neuromagnetic brain responses in a self-paced button press task for both adults and children and have been described in numerous previous publications (e.g. Cheyne et al., 2014; Johnson et al., 2020).

Overall, we conclude that the cluster-based permutation analyses show a differential SAM beamformer effect (p < .05) for all four groups, corresponding to clusters in the observed data at a spatial location corresponding to the known anatomic location of the hand region of sensorimotor cortices in the left hemisphere, contralateral to the right hand used for the button press task. Further, virtual sensor time-spectrograms generated from the cluster centre locations for each group are entirely representative of and in accord with well-known and well-characterised neuromagnetic brain responses in a self-paced button press task.

### Speech data

Group mean statistical analyses of SAM-beamformer images for the speech condition are shown in Figure 2 and the corresponding group mean images are plotted on the fsaverage brain in Figure 3.

**Figure 2.**
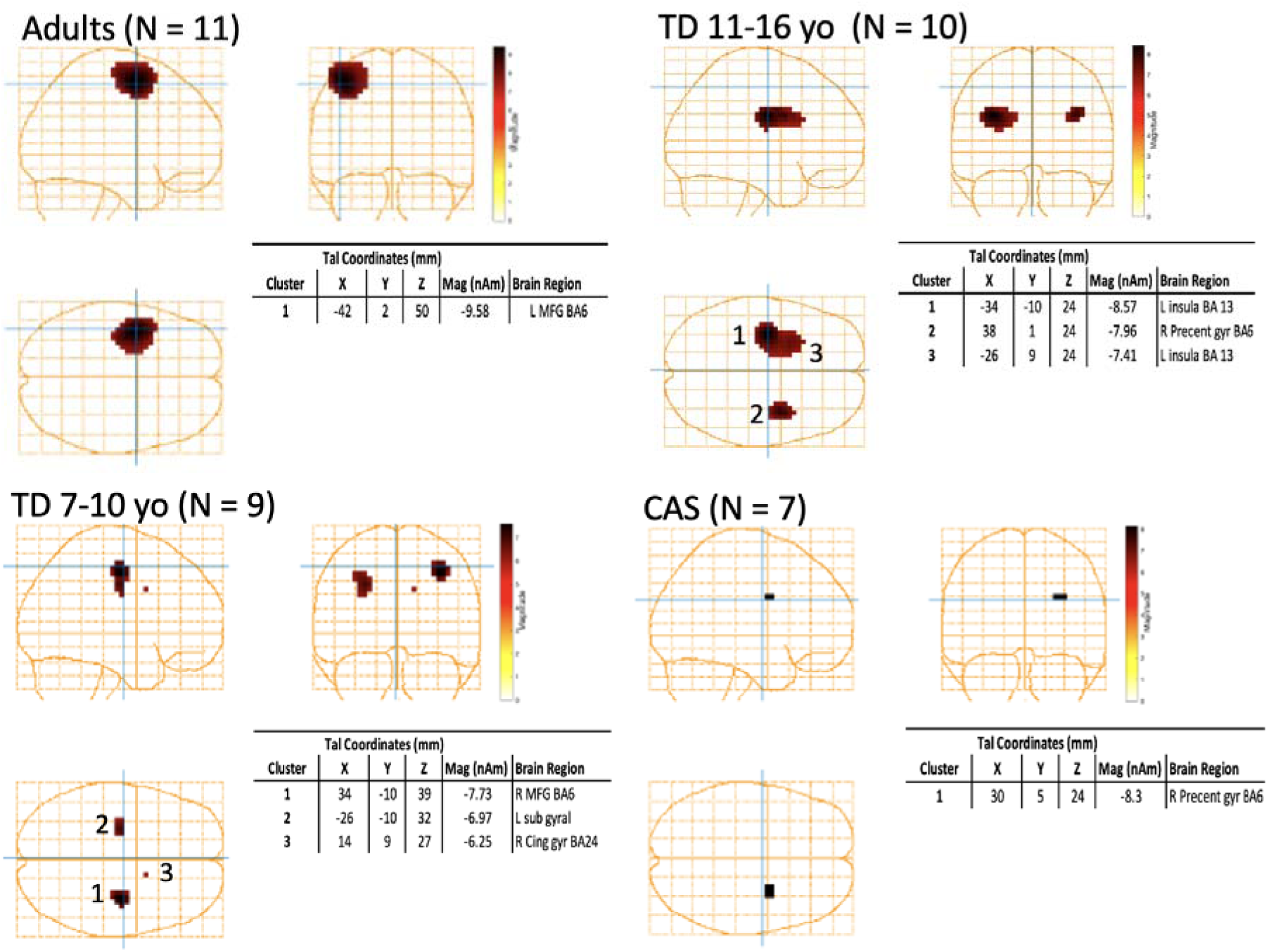
Group statistical maps for speech condition. *Note.* Cluster-based permutation analysis of SAM-beamformer source maps. Cluster alpha = 0.05, 512-1024 permutations, omnibus correction for multiple comparisons. Locations are in Talairach coordinate system. L = left hemisphere, R = right hemisphere, MFG = medial frontal gyrus, BA = Brodmann area. Multiple sources are numbered according to source magnitude.

**Figure 3.**
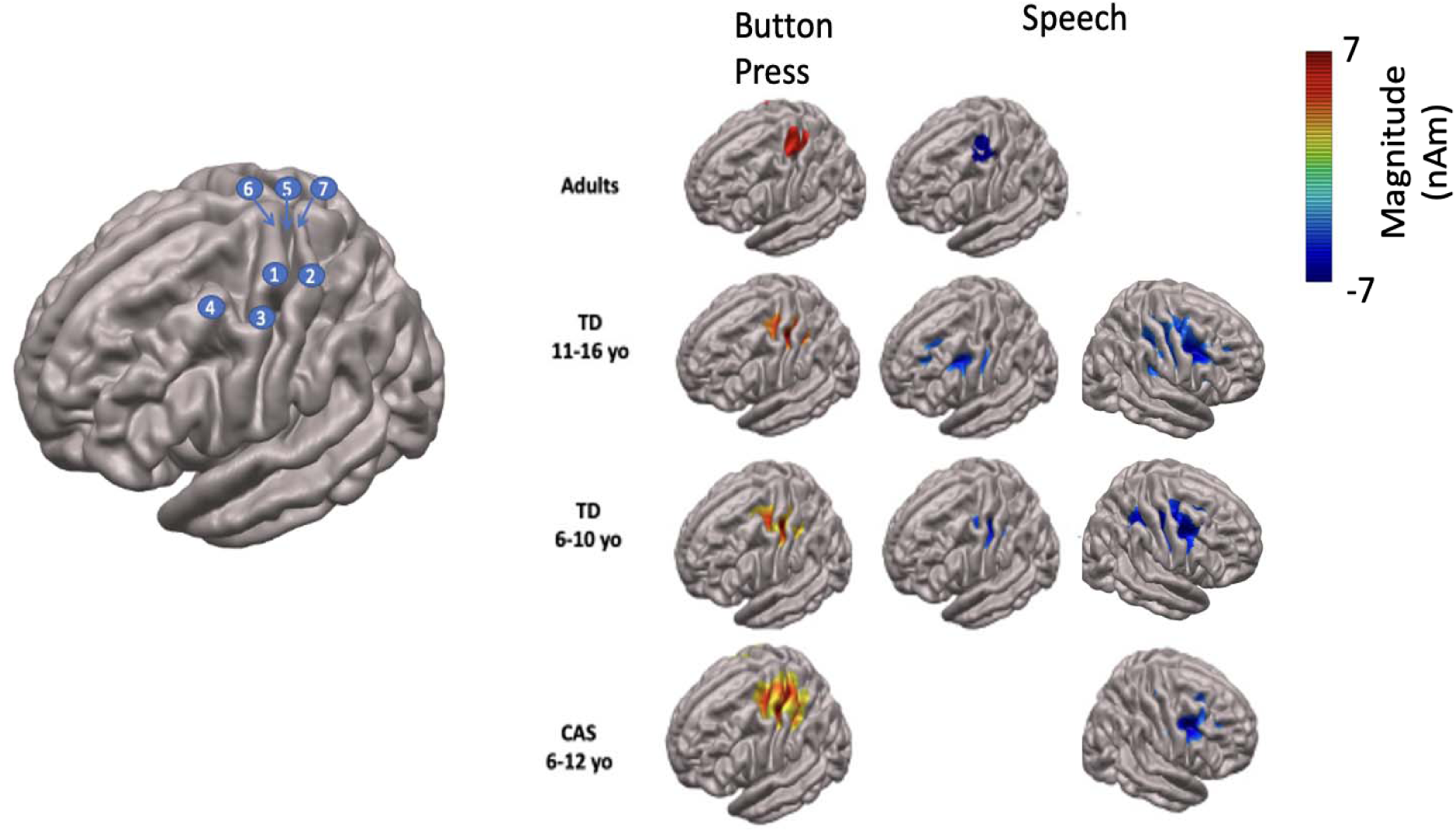
Group mean SAM beamformer maps. *Note.* All plots are shown on fsaverage brain and correspond to the significant left and right hemisphere clusters shown in Table 4 and Figure 2. *Left panel*: Anatomical landmarks. 1 –Hand region of precentral gyrus (hand knob), 2 – Hand region of postcentral gyrus 3 – Middle precentral gyrus, 4 – Middle frontal gyrus, 5 – Rolandic fissure, 6 –Precentral gyrus, 7 – Postcentral gyrus. *Right panels*: Thresholded group mean SAM beamformer maps for button press and speech conditions.

Beginning with the adult group, the statistical maps (Figure 2) show a single extensive cluster in the left hemisphere which is centred on the left medial frontal gyrus (MFG; BA6). Comparison of the group mean button press and speech maps (Figure 3) indicates that the speech related MFG activity map is immediately anterior to the prefrontal gyrus, and that the maps extends posteriorly to encompass a region of the precentral gyrus immediately ventral to the hand area of the precentral gyrus mapped by the button press response.

The virtual sensor time-frequency spectrogram generated from the MFG voxel at the speech cluster centre is shown in Figure 4 (top right panel). The following features can be observed in this plot: (1) Substantial high-frequency broadband noise during the speech trial sets, attributable to muscle activity that is an inevitable artifact of overt speech task; (2) prominent mu/beta band (about 10-25 Hz) desynchronisation that is continuous during the speech trial sets, and also for about 2 seconds prior to the speech trial set onset, reflecting motor preparatory activity that is characteristic of the mu/beta bands (Cheyne, 2013). Relative to the button press response, where the beta band range is about 13-30 Hz, with a mid-frequency about 22-23 Hz, the speech spectrogram displays a lower frequency range (about 10-25 Hz) with a mid-frequency of about 18 Hz. (3) Theta band (circa 3-7 Hz) synchronisation which roughly tracks the time course of the mu/beta desynchronisation. (For aid in interpretation we note that the speech spectrograms are baselined to the first two seconds of the inter-trial set rest period, -5 to -3 sec. This baseline affects the visual appearance of the spectrogram during the immediately preceding time period, where the high-frequency noise and mu/beta-band desynchronisation appear to stop about 1 sec before the end of the speech trial set.)

**Figure 4.**
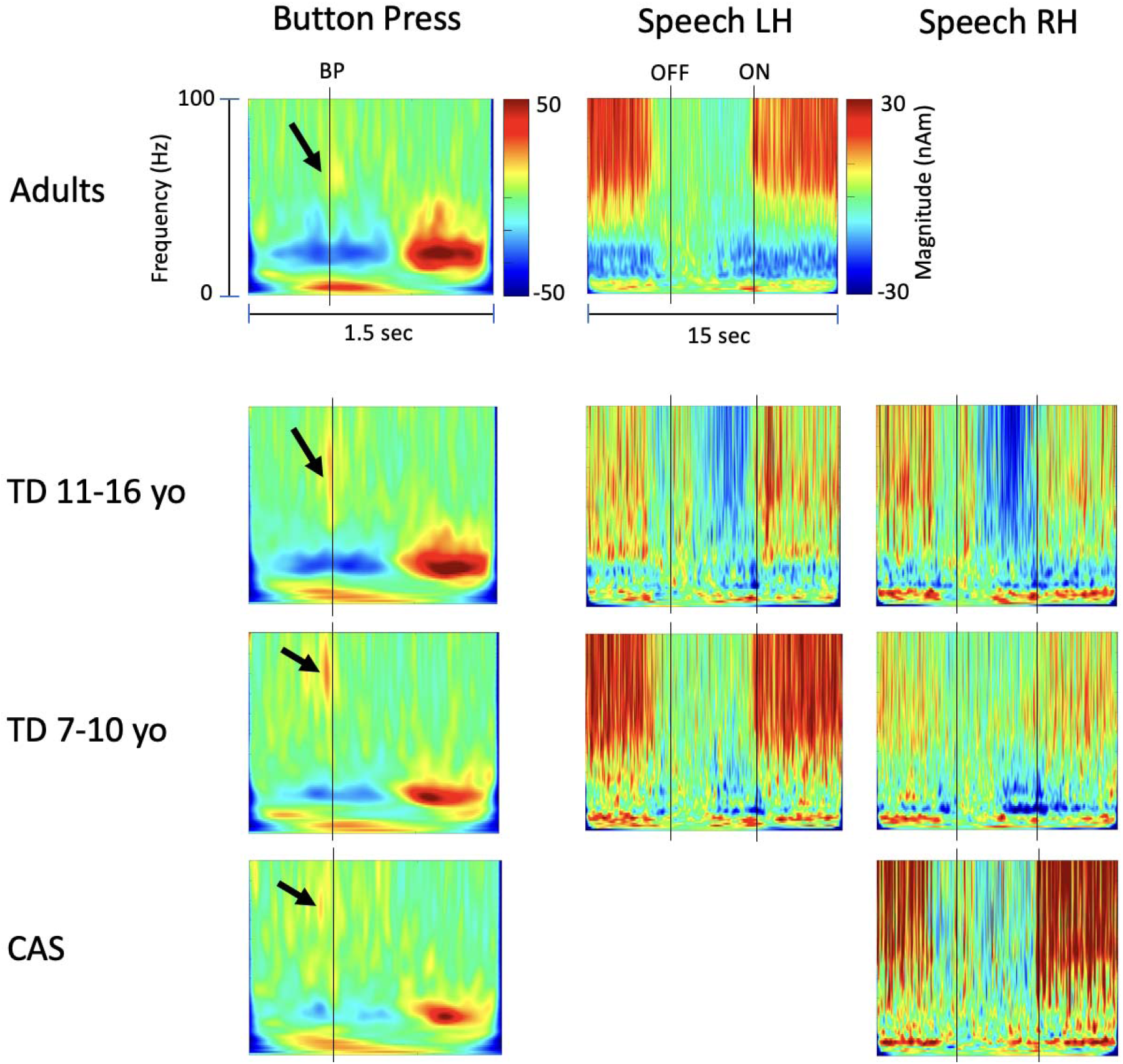
Group virtual sensor time-frequency spectrograms. *Note.* All virtual sensor plots correspond to the centre voxels of significant left and right hemisphere clusters shown in Table 4 and Figure 2. Where more than 2 significant clusters were obtained, virtual sensors for only the first two (largest magnitude) clusters are plotted. Button press data are baselined to the full epoch; speech data are baselined -5 to -3 sec to emphasise beta band desynchronisation. BP = button press onset, OFF = speech trial set offset, ON = speech trial set onset. Arrows indicate gamma band synchronisation around the time of button press onset.

Relative to the adult results, the Figure 2 statistical maps for both groups of TD children show distinctive patterns of significant clusters. First, both TD groups show significant clusters in both cerebral hemispheres, while the adult cluster is restricted to the left hemisphere. Second, the voxel centres of all TD clusters are relatively inferior (*z* = 24-39 mm) to that of the adult group (*z* = 50 mm). Third, the TD clusters appear relatively deeper (absolute *x* of 2 largest magnitude clusters = 26-38 mm) than the adult cluster (absolute *x* = 42).

Group discrepancies in the precise locations of clusters are difficult to interpret from the present data. On the one hand, reasonable between-group agreement of source locations for the button press response supports adequacy of source modelling for all groups: in this case all groups showed good agreement for *x* and *y* coordinates, while the *z* discrepancy is explainable from the larger and more extensive source configuration in the adults. On the other hand, group discrepancies may be expected to arise from at least three sources: first, adult source modelling was based on individual MRI scans, while the child models were derived from template brains; second, the cluster-based analysis indicates both more extensive and stronger (larger magnitude) sources for the adults than for the children, resulting in a group difference in source signal-to-noise ratio that may affect model comparisons; third and perhaps most significantly for this comparison, it is well-known that adaptive beamformers perform suboptimally in the case of correlated bilateral sources, since linear dependencies between the neuronal source timeseries are utilised by these algorithms to minimize the output power (e.g. Kuznetsova et al., 2021).

Overall, these considerations preclude any clear inferences concerning group differences in the precise locations of speech-related sources. Within the stated limitations, the results of the statistical cluster-based analyses suggest the following: for adults, a differential SAM beamformer effect (p < .05) corresponds to a single cluster in the observed data at a spatial location in the precentral gyrus of the left hemisphere immediately inferior to the known anatomic location of the hand region of sensorimotor cortices, and co-extensive with the region of the medial frontal gyrus that is immediately anterior to this middle region of the pre-central gyrus; in contrast, for both TD child groups, the differential SAM beamformer effects (p < .05) correspond to bilateral clusters in both cerebral hemispheres, at relatively lateralised locations on a mid-line (in the sagittal plane) roughly corresponding to the location of the central sulcus/ pre- and post-central gyri in both hemispheres.

Time-frequency spectrograms generated from centre voxel locations in left and right hemisphere clusters for the TD groups are shown in the centre panels of Figure 4. For both groups and both hemispheres the plots show speech-related neural activity that is comparable to that described for the adults above: mu/beta desynchronisation during speech, and for several seconds prior to the onset of the speech trial set. One temporal-spectrographic feature of the children’s data is not observed in the adult data: a distinct low frequency band (circa 7-10 Hz) of synchronisation that roughly follows the time course of the speech-related mu/beta-band desynchronisation. In contrast for the adults, a comparable but less distinct pattern of speech-related synchronisation is seen at lower frequencies confined to the theta band (circa 3-7 Hz).

In contrast to the typically developing children, the cluster analysis of the CAS group shows only a single, small, right hemisphere cluster (Figure 2). The virtual sensor at this location (bottom right panel of Figure 4) shows a speech-related pattern of low frequency (7-10 Hz) synchronisation that is also observed in the TD groups. However, unlike the prominent beta patterns in both adults and TD children, no discernible pattern of speech-related beta-band desynchronisation is observed in this plot.

The lack of speech-related beta-band desynchronisation in the group mean time-frequency spectrogram is entirely representative of the seven individuals within the group. Figure 5 shows that only one individual (S1154) showed speech-related desynchronisation in a narrow mu frequency (circa 10-12 Hz) band. Of the seven, six show speech-related synchronisation in the sub-mu 7-10 Hz band. No individuals show the speech-related beta-band desynchronisation that is prominent in the adults and both TD groups.

**Figure 5.**
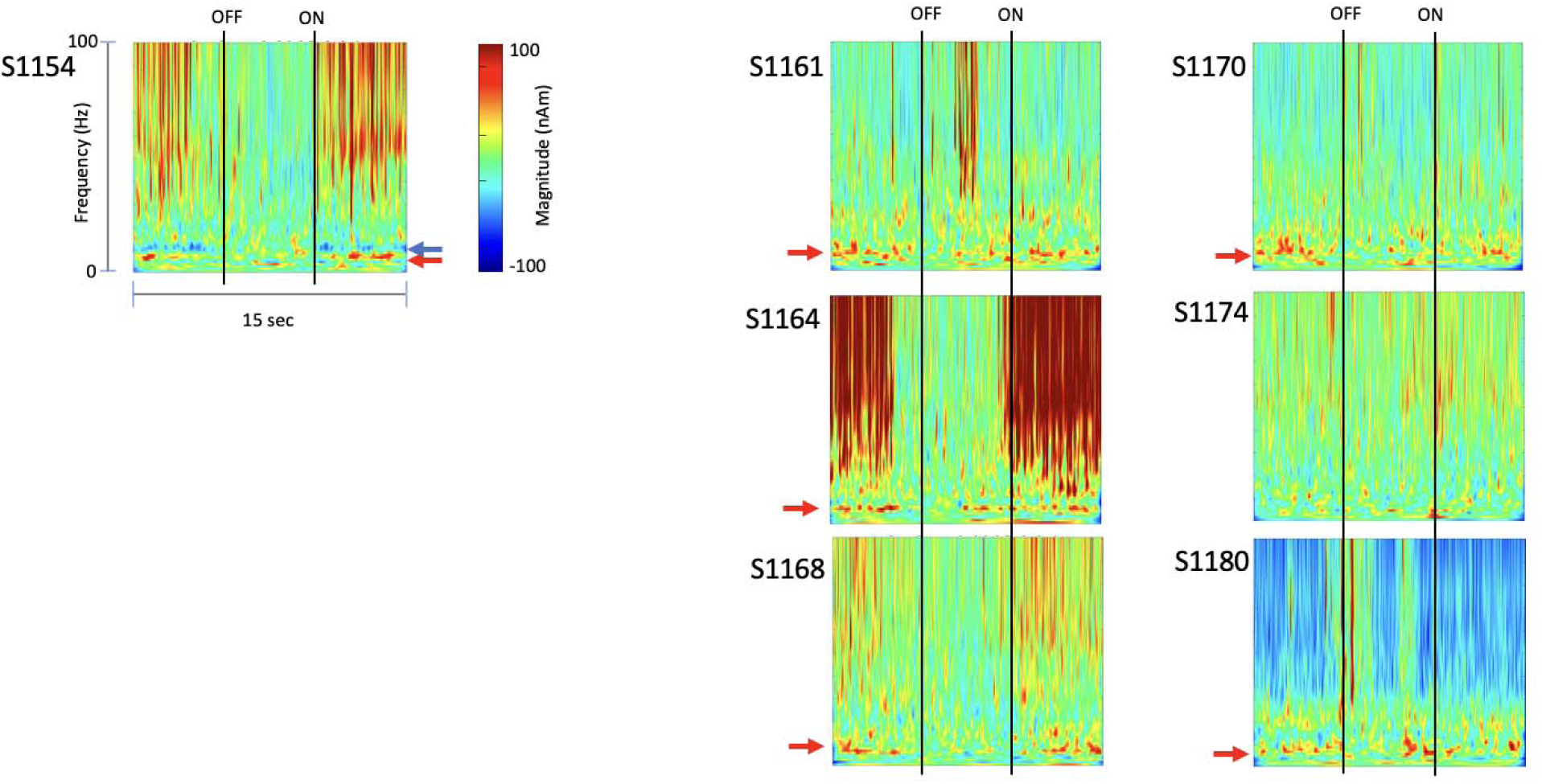
Time-frequency spectrograms for CAS individuals. *Note.* Six of the seven CAS children show speech-related synchronisation in a sub-mu (circa 7-10) Hz frequency band (red arrows). A single participant (S1154) shows speech-related desynchronisation narrowly concentrated within the mu band (circa 10-12 Hz; blue arrow). All plots are generated from the centre voxel of the CAS cluster shown in Figure 2. OFF = speech trial set offset, ON = speech trial set onset.

## Discussion

The present study assessed speech motor cortex activations as elicited by a reiterated non-word speech production task and measured with MEG. Our source-reconstructed results are in good agreement with those reported in the fMRI study by Riecker et al. (2000), indicating quite focal activations in all groups, and in all cases restricted to the central midline axis of the brain corresponding to the location of the peri-Rolandic sensorimotor cortices. This relative focality of speech-elicited responses stands in contrast to the fairly extensive and distributed patterns of brain activity obtained with more commonly employed speech tasks that typically rely on and invoke higher level cognitive and linguistic operations in addition to the phonological, phonetic, and motor operations required at or near the stage of final cerebral output for speech production. In the following, we discuss the following salient observations from these results. First, in adults, our analyses indicate that the locus of significant speech-related clusters can be specified to encompass the left medial frontal gyrus (MFG) and the immediately adjacent region of the pre-central gyrus (middle pre-central gyrus; mPCG) just posterior to the MFG. This middle region of the precentral gyrus is inferior to the hand region of the primary motor strip on the precentral gyrus, and superior to the conventionally defined “ventral speech areas” of the precentral gyrus which encompasses the face/mouth/oral cavities of the familiar sensorimotor homunculus. Second, in contrast, both groups of typically developing children showed bilateral speech-related clusters. Third, CAS children, with a known deficit in the ability to generate proficient reiterated speech, show both atypical right hemisphere lateralisation and significantly reduced speech-related beta band activity relative to both groups of typically developing children.

### The middle precentral gyrus and the dorsal precentral speech area

The functional relevance of the MFG and the midPCG to expressive speech motor control have been highlighted in several recent reviews of lesion, electrocorticographic, and functional neuroimaging evidence (see Gordon et al., 2023; Hickok et al., 2023; Jensen et al., 2023; Silva et al., 2022). In a pivotal study by Chang et al. (2020), a right-handed male subject underwent focal surgical resection of a recurrent grade III astrocytoma located in the left hemisphere’s dorsal premotor cortex, specifically within the posterior middle frontal gyrus. Subsequent to the surgical procedure, the patient exhibited pronounced symptoms indicative of pure apraxia of speech. Comparative analysis was conducted by juxtaposing these findings with publicly available datasets from the Human Connectome Project, revealing that the identified deficits were localized to a specific region, area 55b. This region is recognized as crucial for language function, encompassing both MFG and the midPCG. Furthermore, a recent lesion study conducted by Morshed et al. (2022) retrospectively examined 39 patients diagnosed with gliomas situated within the MFG, necessitating surgical resection as part of their treatment. The results of this study provided further confirmation of the MFG’s pivotal role in language processing. Post-surgery, patients displayed temporary speech hesitancy and dysfluency. These language impairments manifested as speech hesitation, complete speech arrest, naming errors, speech slurring, pronunciation difficulties, and semantic paraphasia. In a recent ECoG (Electrocorticography) study conducted by Castellucci et al. (2022), neural activity was closely observed while participants were actively engaged in conversations. Notably, their findings shed light on the activation patterns within the brain during different phases of speech perception and speech production. They found that Broca’s area and the MFG were both activated when participants were in the process of planning their responses during conversation.

The present results from non-invasive MEG neuroimaging dovetail neatly with an emerging neuroscientific consensus which assigns a central role in motor speech coordination to the middle prefrontal gyrus and adjacent regions of the MFG (Gordon et al., 2023; Jensen et al., 2023; Silva et al., 2022). This new conceptualization is exemplified in the recent “dual motor system” model of expressive speech processing (Hickok et al., 2023), which refers to this region as the “dorsal precentral speech area” (dPCSA) and assigns it a crucial role in speech coordination. On this model, the dorsal precentral speech area is part of a hierarchical control system consisting of the dPCSA, the dorsal laryngeal motor cortex (dLMC) and the posterior portion of the middle frontal gyrus. The dorsal speech system, which shows strong functional connectivity to the auditory cortex, functions to control pitch-related vocalisation, including prosodic aspects of speech (as well as song) and acts primarily on the larynx via the dorsal laryngeal motor cortex. In the dual motor model, the dPCSA works in parallel with a functionally and anatomically distinct ventral control centre (ventral precentral speech area; vPCSA) which corresponds to the more familiar ventral speech areas of the precentral gyrus and Broca’s area in the prefrontal cortex and relies more heavily on functional connections to somatosensory cortex. Accordingly, the dual motor model assigned the vPCSA a function in somatosensory-weighted control of supralaryngeal vocal tract coordination of articulation at the phonetic/syllabic level.

Preferential activation of the dPCSA may well be expected in the context of a continuously reiterated non-word speech task, which is empty of lexical content, repetitive, and is more akin to monotonous song or chant than it is to any form of discrete or extended set of discrete utterances that require continuous precision of individual articulations to ensure intelligibility. On the other hand, it can be expected that reiterated speech would require continuous prosodic control over an extended and continuous period of repetitive utterances: prosodic variations (and continuous corrections) in the form of changes in utterance rate and syllable stress are the most prominent errors that are typically observed in this task.

Relative to the extended and distributed patterns of activations that are typically obtained in lexical speech production tasks, our MEG results in adults show a single, focal, left hemisphere cluster associated with reiterated non-lexical speech. The relative focality and locus in the region of primary sensorimotor cortex agrees well with the fMRI results for reiterated speech reported by Riecker et al. (2000). These authors suggested that overlearned, automatic, and repetitive utterances can be effectively “chunked” at a planning level in a manner that places far fewer demands on neural resources than are required for single “individualised” speech gestures. In addition, the lack of lexical content in the nonword utterances removes the need for any access to higher language centres for syntactic processing or lexical retrieval.

However there are two salient differences between the current MEG results and Riecker et al.’s (2000) fMRI results: Our results are left lateralised and with a dorsal locus corresponding to the dPCSA of the dual motor model, while the fMRI results were bilateral for two of the three nonlexical speech tasks, “ta” and “stra”, and the “pataka” response was left lateralised; the fMRI results also showed a ventral locus on sensorimotor cortex (across all speech tasks), corresponding to the vPCSA. One possibility for the dorsal/ventral difference is suggested by the dual motor model: the Riecker et al. (2000) participants were exposed to loud masking noise caused by gradient switching in the fMRI environment, while the MEG environment is silent with respect to environment noise. Since the dPCSA is dependent on auditory feedback, it is possible that the fMRI participants were forced to rely on somatosensory feedback accessible in the ventral speech control centre. The relative hemispheric lateralisation organisations of the vPCSA and dPCSA are presently unknown (Hickok et al., 2023) so it is possible that a more bilaterally organised dPCSA may also account for the bilateral results in the fMRI study. However, it is also possible that our VCV productions /ipa/ and /api/ are more akin to the CVCVCV structure of “pataka” (which elicited left lateralised fMRI activity) than to the CV and CCCV structures of “ta” and “stra” (which elicited bilateral fMRI activity). These possibilities merit evaluation in future studies.

### Developmental differences in speech motor control

Our results in adults and typically developing children are entirely in accord with previous neuroimaging work indicating a developmental shift from bilateral hemispheric control of speech and language in childhood to left hemisphere dominance in adulthood (see Holland et al., 2007, for a review of fMRI results). Two previous MEG studies (Kadis et al., 2011; Ressel, 2008) have also reported that speech-related MEG responses are bilateral in young children and show an age-related increase in left hemisphere dominance.

Relative to the adult results, the typically developing groups also showed speech-elicited clusters that were deeper and more ventral. Group discrepancies in the precise locations of clusters are difficult to interpret from the present data. On the one hand, reasonable between-group agreement of source locations for the button press response supports adequacy of source modelling for all groups: in this case all groups showed good agreement for *x* and *y* coordinates, while the *z* discrepancy is explainable from the larger and more extensive source configuration in the adults. On the other hand, group discrepancies may be expected to arise from at least three sources: first, adult source modelling was based on individual MRI scans, while the child models were derived from template brains; second, the cluster-based analysis indicates both more extensive and stronger (larger magnitude) sources for the adults than for the children, resulting in a group difference in source signal-to-noise ratio that may affect model comparisons; and third, it is well-known that adaptive beamformers perform suboptimally in the case of correlated bilateral sources, since linear dependencies between the neuronal source timeseries are utilised by these algorithms to minimize the output power (e.g., Kuznetsova et al., 2021).

Overall, these considerations preclude any clear inferences concerning adult/TD group differences in the precise locations of speech-related sources. Within the stated limitations, the results of the statistical cluster-based analyses suggest the following: for adults, a differential SAM beamformer effect (p < .05) corresponds to a single cluster in the observed data at a spatial location in the precentral gyrus of the left hemisphere immediately inferior to the known anatomic location of the hand region of sensorimotor cortices, and co-extensive with the region of the medial frontal gyrus that is immediately anterior to this middle region of the pre-central gyrus; in contrast, for both TD child groups, the differential SAM beamformer effects (p < .05) correspond to bilateral clusters in both cerebral hemispheres, at relatively lateralised locations on a mid-line (in the sagittal plane) roughly corresponding to the location of the central sulcus/ pre- and post-central gyri in both hemispheres.

### Childhood Apraxia of Speech

At the present time, the neuroimaging and neurophysiological evidence base for CAS is notably small and the putative central deficits remain highly underspecified. Although there is a consensus that the deficits of CAS have a central basis (Morgan et al., 2016; Liégeois et al., 2014; Liégeois & Morgan, 2012) there is little understanding of the precise brain abnormalities underlying the speech motor deficits that characterise CAS; and how these brain abnormalities are distinct from or overlap with those associated with other classes of speech difficulties. Moreover, there is evidence from a few magnetic resonance imaging (MRI) studies investigating neural activity in children with idiopathic CAS, however with inconsistent findings. Unilateral brain abnormalities have been reported either in left (Fiori et al.’s 2016; Kadis et al. 2014) or in the right hemisphere (Preston et al., 2014), in contrast to the bilateral brain abnormalities that were a prominent feature of the FOXP2-related CAS patients’ studies by Liégeois & Morgan (2012). White matter reductions in temporal regions of the left hemisphere as well as in the inferior frontal gyrus were reported by Fiori et al. (2016) who measured white matter connectivity in children with idiopathic CAS using whole brain tractography. Kadis et al. (2014) reported significantly increased volume of the left supramarginal gyrus in children with idiopathic CAS after a speech and language intervention. However, the volume effect was observed only at the group level; they found no significant correlation between these brain changes and behavioural assessments of individual changes in speech production. Contradictory results were reported by an electroencephalography (EEG) study of children with idiopathic CAS (Preston et al., 2014). They reported unilateral dysfunctions, specifically a reduced amplitude of EEG responses in the right hemisphere when children with CAS processed more complex words.

To date, the most comprehensive research findings regarding the neurobiology of CAS have been carried out in neurogenetic cases with mutations of the *FOXP2* gene. Neuroimaging of 26 affected KE family members has revealed structural and functional anomalies of cortical and subcortical brain networks (Liégeois et al., 2019; Liégeois et al., 2011). Abnormal increases or decreases of grey matter volume were found in the caudate nucleus, putamen (Liégeois et al., 2011; Liégeois et al., 2003; Watkins et al., 2002), cerebellum (Belton’s et al., 2003); globus pallidus (Liégeois et al., 2016); motor cortex and inferior frontal gyrus (Watkins et al., 2002; Belton’s et al., 2003). The basal ganglia have been strongly implicated in language processing (Argyropoulos et al. 2013), and bilateral abnormalities in the basal ganglia have been reported in all KE family members with a FOXP2 gene mutation (Watkins et al., 2002; Belton et al., 2003; Liégeois et al., 2003; Liégeois et al., 2016). However, contradictory results have been obtained for CAS participants from another family whose first- and second-generation members were diagnosed with the literacy and language impairments features of CAS (the specific gene or genes have not been identified). In this family, participants did not show any significant basal ganglia deficits (Liégeois et al., 2019). Further, the FOXP2 mutation is not a diagnostic criterion for CAS (Preston et al., 2014) and the FOXP2 neuroimaging results cannot be extrapolated to all cases with CAS.

Variations in methodological approaches, speech tasks and the ages of participants make comparisons difficult across the small number of MRI and fMRI studies of neurogenetic CAS. Thus, different qualitative imaging techniques and analyses have been used, including voxel-based morphometry, functional MRI, magnetic resonance diffusion-weighted imaging and fibre-tractography. Further, all of the cited studies have employed group-level analyses, and none are currently capable of identifying CAS brain lesions at an individual level (Kurth et al., 2018; Morgan et al., 2016; Preston et al., 2014; Vargha-Khadem et al., 2005). Moreover, a variety of speech tasks tapping different aspects of spoken language have been used. Liégeois and colleagues (2019; 2003) and Belton et al., (2003) used a word repetition task paradigm to investigate activation differences during speech production in the affected members and in controls during functional and structural brain scanning. In contrast, the experimental tasks used by Liégeois et al. (2011; 2016) consisted of repetition of nonwords during functional MRI scanning. With regards to age, different studies have recruited KE family members of the second and third generation within different age groups (Belton et al., 2003; Watkins et al., 2002; Liégeois et al., 2016). In Liégeois et al.’s (2011) study, participants were adults, but their exact ages were not specified.

The results of the present study provide an important step forward in our understanding of the neural deficits that are likely to underlie the speech problems of CAS. Relative to both younger and older children with typical speech development, the CAS group showed no evidence of speech-related activity in the left hemisphere: the differential SAM beamformer effect (p < .05) corresponded to a single cluster in the observed data at a relatively lateralised location on a mid-line (in the sagittal plane) roughly corresponding to the location of the central sulcus/ pre- and post-central gyri in the right hemisphere. Virtual sensor time frequency plots generated from this location showed that as a group, and also for every individual, there was no evidence for the speech-related beta-band desynchronisation that is a prominent temporal-spectral feature of bilateral hemispheric clusters in both groups of TD children and the left hemispheric cluster in adults.

The major weakness of this CAS data is the relatively small group size (N=7), which is fairly typical of published studies of this rare disorder. Offsetting this weakness are a number of important methodological and analytical strengths: The CAS phenotype was rigorously characterised with a comprehensive battery of clinical tests and diagnosed independently on the basis of overt speech behaviours by two experienced speech-language pathologists; the locus of the small CAS group right hemisphere cluster is in accord with the right hemisphere clusters of both groups of TD children; and the neurophysiological temporal-spectral profiles of all seven individuals were in agreement in the absence of speech-related beta-band activity that is a prominent feature of both TD groups and the adults. Therefore, despite the small sample size, group beta-band magnitude differences were statistically significant for both the CAS-TD younger group comparison and the CAS-TD older group comparison. The large effect size strongly suggests that the speech-related beta motor rhythm is an important neurophysiological marker of both typical and apraxic speech development.

## Conclusions

The results of the present study indicate that reiterated nonword speech elicits patterns of brain activity that are more restricted and focal than those obtained with the more individualized and lexical speech production tasks in common clinical and experiment use. Since by their nature MEG source reconstruction techniques favour focal configurations, the reiterated speech task may provide a more precise and informative mapping of expressive language function than existing MEG speech production protocols where outcomes remain largely limited to a basic index of hemispheric laterality. Our results are in agreement with previous studies showing bilateral organisation of speech in children developing to left hemispheric lateralisation in adulthood. Finally, our results indicate that the speech-related beta frequency motor rhythms is a robust neurophysiological marker of typical and atypical speech development.

## Notes

### Competing Interest Statement

The authors have declared no competing interest.

### Summary of Updates

Minnor changes in formatting

